# TERRA R-loops connect and protect sister telomeres in mitosis

**DOI:** 10.1101/2023.05.31.543102

**Authors:** Samantha Sze, Amit Bhardwaj, Priyanka Fnu, Kameron Azarm, Rachel Mund, Katherine Ring, Susan Smith

## Abstract

Resolution of cohesion between sister telomeres in human cells depends on TRF1-mediated recruitment of the polyADP-ribosyltransferase, tankyrase to telomeres. In cells where tankyrase is deleted or the tankyrase binding site in TRF1 is mutated, sister telomeres remain cohered in mitosis. Human aged cells and ALT cancer cells naturally exhibit persistent telomere cohesion due to shortened telomeres that do not recruit sufficient TRF1/tankyrase for resolution. Persistent cohesion plays a protective role, but the mechanism by which sister telomeres remain cohered is not well understood. Here we show that telomere repeat containing RNA (TERRA) holds sister telomeres together through RNA-DNA hybrid (R-loop) structures. We show that a tankyrase-interacting partner, the RNA-binding protein C19orf43 is required for resolution of telomere cohesion and for repression of TERRA R-loops. Depletion of C19orf43 led to persistent telomere cohesion and an increase in TERRA R-loops. Overexpression of RNaseH1 counteracted persistent cohesion in C19orf43-depleted cells, as well as in aged and ALT cells. In fact, treatment of cohered telomeres in mitotic cells with RNaseH1 in situ, was sufficient to resolve sister telomere cohesion, confirming that RNA-DNA hybrids hold sister telomeres together. Consistent with a protective role for persistent telomere cohesion, depletion of C19orf43 in aged cells reduced DNA damage and significantly delayed replicative senescence. We propose that the inherent inability of shortened telomeres to recruit R-loop repressing machinery permits a controlled onset of senescence.

## Introduction

Human telomeres consist of tandem arrays of double-stranded TTAGGG repeats, telomere repeat containing RNA (TERRA), and the shelterin complex. Shelterin comprises six subunits, two of which, TRF1 and TRF2, bind directly and specifically to the double-stranded TTAGGG repeats(de Lange, 2005). Shelterin and associated factors collaborate to: protect chromosome ends from being viewed as double strand breaks; regulate telomere length and replication; and control maintenance and resolution of telomere cohesion(Azarm and Smith, 2020; de Lange, 2018; Lim and Cech, 2021).

Resolution of cohesion between sister telomeres prior to mitosis requires the polyADP- ribosyltransferases, tankyrase 1 and tankyrase 2. A distinguishing feature of tankyrases is their ankyrin repeat domains, which can serve as platforms for multiple binding proteins through a consensus tankyrase binding site (TBS), RXX(G/P/A)XG (Azarm and Smith, 2020; Eisemann et al., 2016; Guettler et al., 2011; Sbodio and Chi, 2002; Seimiya and Smith, 2002). Tankyrases localize to telomeres by binding TRF1 in late S/G2 to resolve cohesion(Bisht et al., 2013; Bisht et al., 2012). In cells depleted of tankyrase 1 and/or 2 or where the TBS in endogenous TRF1 is mutated, sister telomeres remain cohered in mitosis despite resolution of arms and centromeres(Azarm et al., 2020; Bhardwaj et al., 2017; Canudas et al., 2007; Dynek and Smith, 2004). Cells with persistent telomere cohesion undergo a prolonged anaphase, but ultimately telomeres resolve and cells exit mitosis(Kim and Smith, 2014).

Despite the positive actions of shelterin and associated factors, telomere function becomes compromised in normal human cells. Due to the end-replication problem and nucleolytic processing, telomeres shorten following each round of DNA replication(Huffman et al., 2000; Wu et al., 2012). This shortening can be counteracted by telomerase, which adds telomere repeats de novo to chromosomes ends(Greider and Blackburn, 1985; Greider and Blackburn, 1987). However, telomerase is repressed in the human soma(Wright et al., 1996). Thus, as cells age, the resulting shortened telomeres are unable to recruit sufficient shelterin to protect the ends, leading to a persistent DNA damage response that signals replicative senescence(d’Adda di Fagagna et al., 2003). Cancer cells must acquire a telomere maintenance mechanism to override senescence; most up-regulate telomerase, but 10-15% activate a recombination-based, alternative lengthening of telomeres (ALT) mechanism(Bryan et al., 1997; Bryan et al., 1995). As a result of the recombination, ALT cells have very long heterogenous telomeres, but a fraction of their telomeres are critically short, like aged cells(Henson et al., 2002).

Interestingly, ALT cancer cells and normal aged cells have another feature in common: persistent telomere cohesion. Thus, similar to tankyrase-depleted or TRF1-mutated cells, telomeres of aged and ALT cells are persistently cohered into mitosis(Kim and Smith, 2014; Ofir et al., 2002; Ramamoorthy and Smith, 2015; Yalon et al., 2004). Persistent cohesion is a direct consequence of telomere shortening; it can be rescued by introduction of telomerase into aged or ALT cells(Azarm et al., 2020; Yalon et al., 2004) and conversely, be induced by long-term inhibition of telomerase in telomerase positive cancer cells (Azarm et al., 2020). Short telomeres in aged and ALT cells do not recruit sufficient TRF1/tankyrase to resolve cohesion. Forced resolution of cohesion by overexpression of TRF1 results in subtelomere recombination with non-sisters, DNA damage, and a senescent-like growth arrest. This indicates that persistent cohesion is a naturally occurring protective state at short telomeres (Azarm et al., 2020; Ramamoorthy and Smith, 2015).

In addition to shelterin and associated factors, telomeres are bound by the telomere repeat-containing RNA TERRA(Azzalin et al., 2007; Luke et al., 2008; Schoeftner and Blasco, 2009). TERRA is transcribed by RNA polymerase II from subtelomeric promoters into the telomere tract toward the end of chromosomes using the telomeric C-rich strand as the template(Azzalin and Lingner, 2015; Azzalin et al., 2007; Feretzaki et al., 2019; Nergadze et al., 2009; Porro et al., 2014). TERRA is retained at telomeres through association with telomeric proteins or by base-pairing with telomeric DNA to form R-loop structures consisting of an RNA/DNA hybrid and a displaced DNA strand(Fernandes et al., 2021; Toubiana and Selig, 2018). R-loops can form during transcription in cis or post transcriptionally in trans(Feretzaki et al., 2020). TERRA is implicated in telomere length regulation by telomerase, telomere replication, and the telomeric DNA damage response(Bettin et al., 2019; Lalonde and Chartrand, 2020). TERRA R-loops also appear to play important roles in telomerase negative cells. In ALT cells TERRA levels are high and TERRA R-loops are regulated by the RNA endonuclease RNaseH1 to control recombination(Arora et al., 2014; Silva et al., 2021). In telomerase negative budding yeast, TERRA R-loops are regulated by RNaseH1 and 2; R-loops accumulate at short telomeres and slow the rate of replicative senescence(Balk et al., 2013; Graf et al., 2017). TERRA and TERRA R-loops have been detected in normal human cells and in ICF Syndrome patient cells but the impact on replicative senescence has not been determined(Sagie et al., 2017; Yehezkel et al., 2008).

Compelled by the similarities between the role of TERRA RNA-DNA hybrids (in ALT cells and in telomerase negative budding yeast) and persistent telomere cohesion (in ALT and aged human cells), we investigated a role for TERRA R-loops in persistent telomere cohesion. We show that overexpression of RNaseH1 in cells or incubation of purified RNaseH1 in situ can resolve sister telomere cohesion. We further identify a tankyrase interacting RNA-binding protein C19orf43 whose depletion results in persistent telomere cohesion, increased TERRA R-loops, and a delayed replicative senescence, indicating a beneficial role for R-loops at aged human telomeres.

## Results

### C19orf43 is a tankyrase binding protein that is required for resolution of telomere cohesion

Tankyrase is recruited to telomeres by TRF1 to resolve cohesion. Its ankyrin repeat domain allows binding to multiple TBS-containing partners at once, imparting a scaffolding capacity, which we reasoned could play a role in resolution of cohesion. Two proteomic screens identified the uncharacterized protein C19orf43, as a potential tankyrase binding protein(Bhardwaj et al., 2017; Li et al., 2017). C19orf43 was also identified as a component of a partially purified nuclear extract and shown to have exoribonuclease activity in vitro(Xie et al., 2017). It was named human telomerase RNA interacting RNase (hTRIR). However, no evidence was presented for a specific association with hTR. It was shown to have 5’ and 3’ exoribonuclease activity in vitro across a wide range of substrates, including hTR. A subsequent study identified C19orf43 as a Protein Phosphatase 4 (PP4) Interacting Protein (IP) and renamed it PP4IP(Park et al., 2019). In our study, motivated by its potential interaction with tankyrase and RNA, we queried the role of C19orf43 in telomere cohesion.

C19orf43 has two potential TBSs, but no other obvious motifs (Fig. 1A). To validate the interaction between C19orf43 and tankyrase, HEK93T cells extracts were immunoprecipitated with anti-TNKS or control IgG. As shown in Fig. 1B, endogenous C19orf43, specifically co-immunoprecipitated with endogenous TNKS. We generated stable cell lines depleted for C19orf43 using two different C19orf43 shRNA-expressing lentiviruses (#1 and #4). Immunoblot analysis showed efficient depletion with both shRNAs (Fig. 1C). We then asked if the potential TNKS-binding sites in C19orf43 were required for interaction with tankyrase. We generated a triple Flag epitope-tagged allele of C19orf43 and mutated the essential R in each of the binding sites to generate mutants R7G and R106G. Vector control, C19orf43.WT, and the C19orf43 mutant plasmids R7G and R106G were each transfected into the HEK293T C19orf43 shRNA#1 cell line. The shRNA#1 targets the 3’untranslated region of C19orf43 and therefore does not affect the transfected constructs as they contain only C19orf43 coding sequence. Transfected cell extracts were immunoprecipitated with anti-Flag antibody. As shown in Fig. 1D, FlagC19orf43.WT and R106G, but not R7G, co-immunoprecipitated endogenous tankyrase, indicating that the amino terminal (but not the internal) TBS is required for C19orf43 interaction with tankyrase.

**Figure 1.**
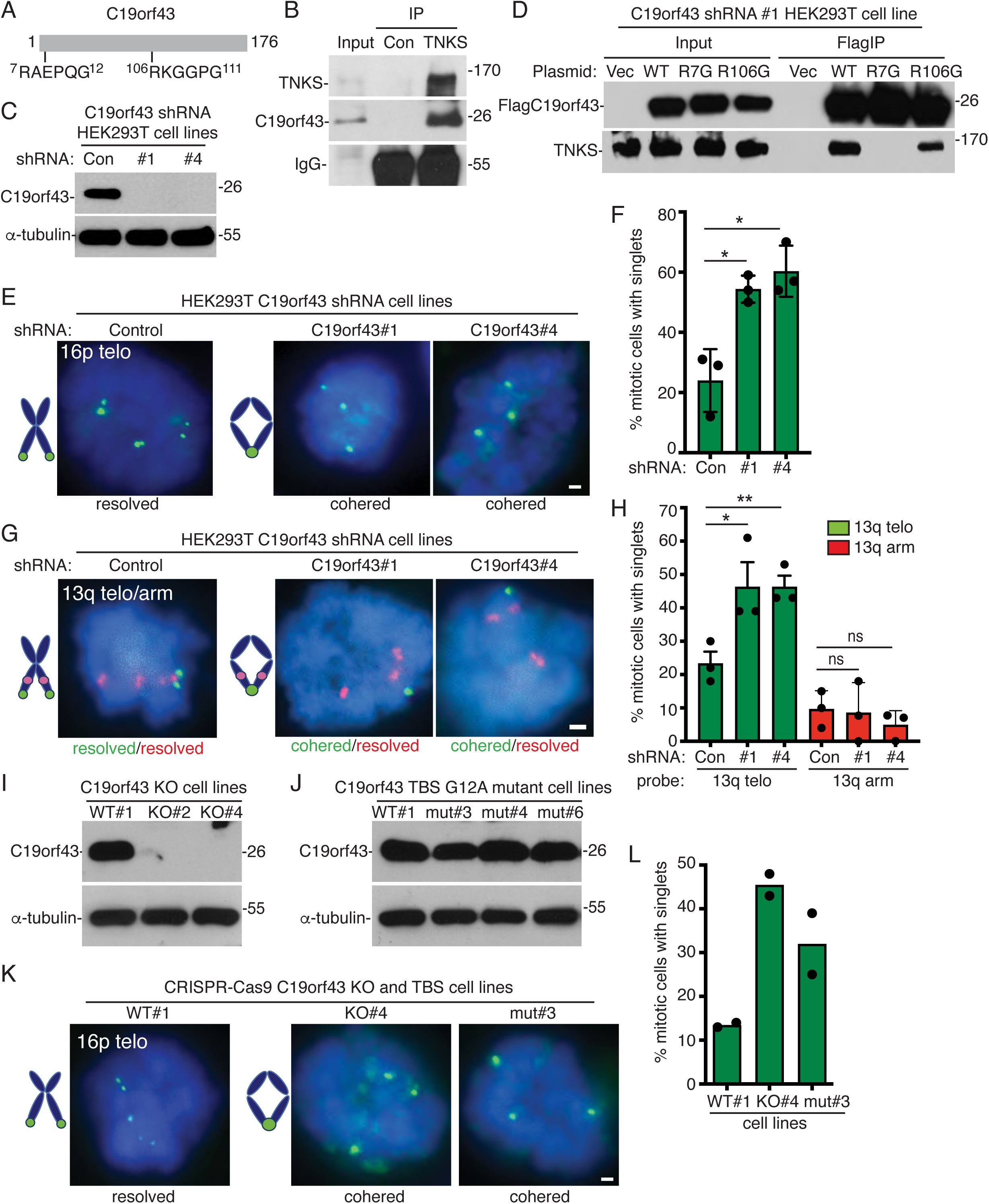
C19orf43 is a tankyrase binding protein that is required for resolution of telomere cohesion. (A) Schematic of C19orf43 with tankyrase binding sites (TBS) indicated. (B) Immunoblot analysis of HEK293T cell extracts immunoprecipitated (IP) with control or anti-TNKS antibody. (C) Immunoblot analysis of cell extracts from HEK293T cells stably expressing control, or C19orf43 shRNAs #1 or #4. (D) Immunoblot analysis of cell extracts from C19orf43 shRNAs #1 HEK293T cells transfected with vector or C19orf43 WT, R7G, or R106G and immunoprecipitated with anti-Flag antibody. (E) FISH analysis of HEK293T control or C19orf43 shRNA (#1 or #4) mitotic cells using a 16p subtelo probe (green). (F) Quantification of the frequency of mitotic cells with cohered telomeres. Three independent experiments (n=37-69 cells each) ± SEM. *p :: 0.05, Student’s unpaired t test. (G) FISH analysis of HEK293T control or C19orf43 shRNA (#1 or #4) mitotic cells with a dual 13q subtelo (green)/arm (red) probe. (H) Quantification of the frequency of mitotic cells with cohered telomeres and arms. Three independent experiments (n=26-29 cells each) ± SEM. *p :: 0.05, **p :: 0.01, Student’s unpaired t test, ns; not significant. (I) Immunoblot analysis of cell extracts from CRISPR/Cas9 generated HEK293T knockout (KO) cell lines #2 and #4. (J) Immunoblot analysis of cell extracts from CRISPR/Cas9 generated HEK293T mutant (mut) cell lines #3, #4, and #6. (K) FISH analysis of WT, KO#4, or mut#3 HEK293T mitotic cells using a 16p subtelo probe (green). (L) Quantification of the frequency of mitotic cells with cohered telomeres. Two independent experiments (n=44-56 cells each). (E, G, K,) DNA was stained with DAPI (blue). Scale bars represent 2 μm.

To determine if C19orf43 is required for resolution of cohesion, we analyzed the shRNA cell lines using fluorescent in situ hybridization (FISH). Mitotic cells were isolated from asynchronous cultures by shake-off, fixed, cytospun onto coverslips, and probed with the subtelomere-specific chromosome probe 16p. As shown in Fig. 1E, F, in control mitotic cells, telomeres appeared as doublets (resolved). However, in C19orf43-depleted cell lines #1 and #4, telomeres appeared as singlets (cohered) in mitosis, indicating persistent telomere cohesion. To determine if the persistent cohesion was specific to telomeres, we performed FISH using a dual telomere/arm 13q probe. Note, HEK293T cells have a triple locus for the 13q arm, but due to a subtelomere deletion, only a single locus for the 13q telomere. FISH analysis showed that the telomere, but not the arms, exhibited persistent telomere cohesion in C19orf43 depleted cells (Fig. 1G, H). Thus, C19orf43 is required specifically for telomere cohesion.

We further validated the role of C19orf43 in persistent telomere cohesion using CRISPR/Cas9 to generate C19orf43 HEK293T knockout cell lines: KO#2 and KO#4 (Fig. 1I) and knock-in cell lines where the essential G in the amino terminal TBS of the endogenous C19orf43 gene was mutated to A to generate C19orf43 mutant G12A cell lines: mut#3, mut#4, and mut#6 (Fig. 1J). FISH analysis showed that both the C19orf43 KO (#4) and the C19orf43.G12A mutant (#3) cell lines exhibited persistent telomere cohesion (Fig. 1K, L).

### C19orf43 rescues persistent telomere cohesion aged and ALT cells

To confirm that C19orf43 itself promotes resolution of telomere cohesion and that its N-terminal TBS is required, we transfected Vector or C19orf43 (WT, R7G, or R106G) into the C19orf43-depleted (shRNA#1) cell line, performed immunoblot analysis (Fig. 2A), and measured telomere cohesion by FISH analysis. As shown in Fig. 2B, C, FlagC19orf43.WT and R106G (but not R7G) rescued persistent telomere cohesion, indicating that C19orf43 (and its amino terminal TBS) is required for resolution of cohesion. We further demonstrated that in cells deleted of C19orf43 (KO#4 cells) reintroduction of C19orf43.WT rescued the persistent cohesion (Fig. 2D, E).

**Figure 2.**
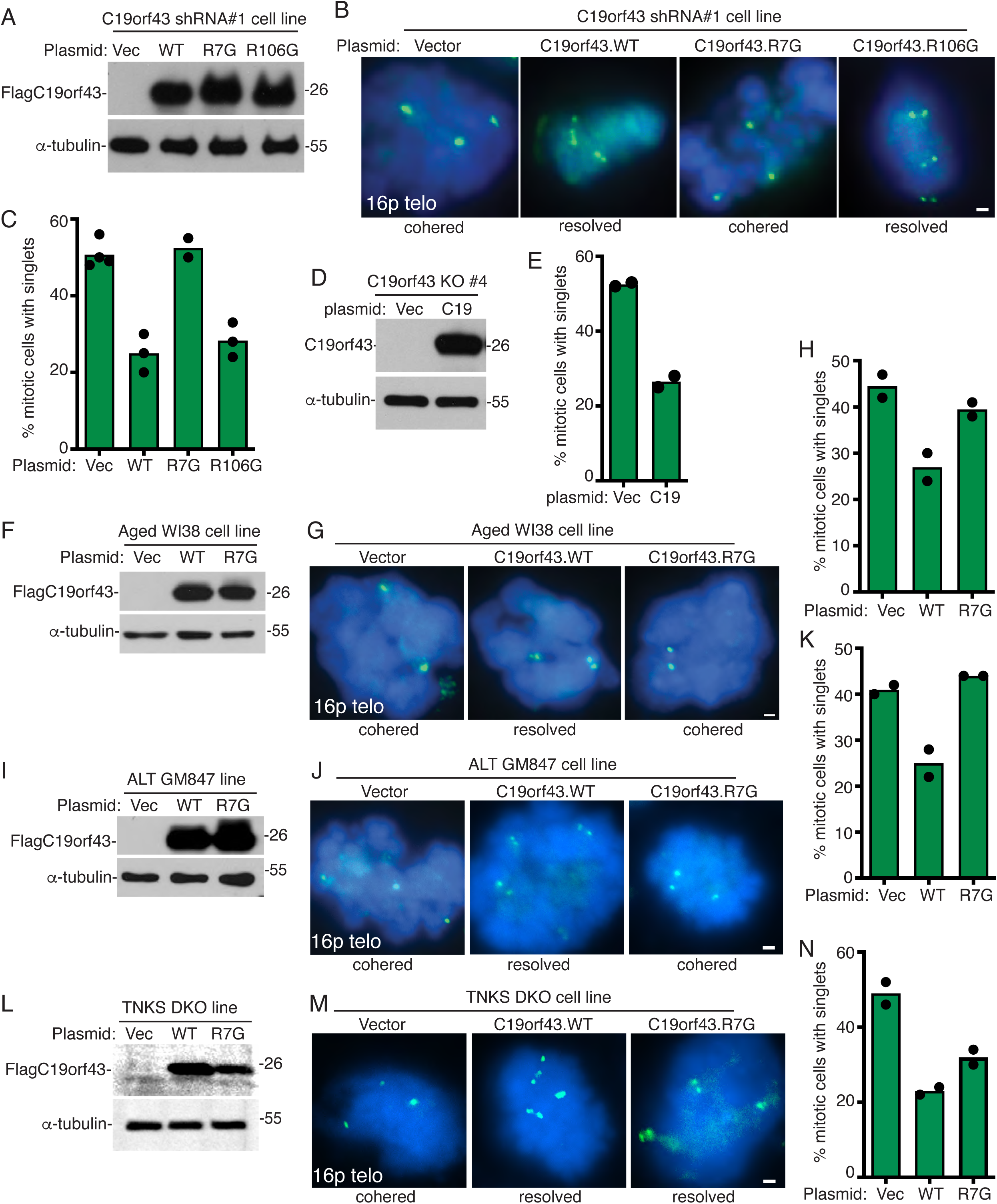
C19orf43 overexpression rescues persistent telomere cohesion in C19orf43 depleted, aged WI38, and ALT cells. (A) Immunoblot analysis of cell extracts from C19orf43 shRNA#1 HEK293T cells transfected with vector or C19orf43 (WT, R7G, or R106G). (B) FISH analysis of vector or C19orf43 (WT, R7G or R106G) transfected C19orf43 shRNA#1 HEK293T mitotic cells with a 16p subtelo probe (green). (C) Quantification of the frequency of mitotic cells with cohered telomeres. Two to four independent experiments (n=27-68 cells each). (D) Immunoblot analysis of cell extracts from C19orf43 KO#4 cells transfected with a vector control or C19orf43 WT. (E) Quantification of FISH analysis showing the frequency of mitotic cells with cohered telomeres. Two independent experiments (n=51-56 cells each). (F) Immunoblot analysis of cell extracts from vector or C19orf43 (WT or R7G) transfected aged WI38 cells. (G) FISH analysis of vector or C19orf43 (WT or R7G) transfected aged WI38 mitotic cells using a 16p subtelo probe (green). (H) Quantification of the frequency of mitotic cells with cohered telomeres. Two independent experiments (n=36-50 cells each). (I) Immunoblot analysis of cell extracts from vector or C19orf43 (WT or R7G) transfected ALT GM847 cells. (J) FISH analysis of vector or C19orf43 (WT or R7G) transfected ALT GM847 mitotic cells using a 16p subtelo probe (green). (K) Quantification of the frequency of mitotic cells with cohered telomeres. Two independent experiments (n=50 cells each). (L) Immunoblot analysis of cell extracts from vector or C19orf43 (WT or R7G) transfected TNKS DKO HEK293T cells. (M) FISH analysis of vector or C19orf43 (WT or R7G) transfected TNKS DKO HEK293T mitotic cells using a 16p subtelo probe (green). (N) Quantification of the frequency of mitotic cells with cohered telomeres. Two independent experiments (n=50 cells each). (B, G, J, M) DNA was stained with DAPI (blue). Scale bars represent 2 μm.

Next, we asked if C19orf43 plays a role in the persistent telomere cohesion that is a naturally occurring feature of aged and ALT cells. We transfected presenescent WI38 cells with a Vector control or C19orf43 (WT or R7G), performed immunoblot (Fig. 2F), and FISH analysis. As shown in Fig. 2G, H, C19orf43.WT (but not R7G) reduced persistent telomere cohesion. Similar results were obtained with transfection into ALT GM847 cells; C19orf43.WT (but not R7G) reduced persistent telomere cohesion (Fig. 2I, J, K). Thus, C19orf43 is required for resolution of telomere cohesion and its overexpression in aged or ALT cells can rescue the persistent telomere cohesion phenotype, dependent on its ability to interact with tankyrase. Finally, we asked if C19orf43-mediated resolution of telomere cohesion was fully dependent on tankyrase for its resolving activity. We transfected Vector or C19orf43 (WT or R7G) into TNKS DKO cells, performed immunoblot (Fig. 2L), and measured telomere cohesion. As shown in Fig. 2M, N, C19orf43 rescued the persistent cohesion, but here (unlike in the cell types above that express tankyrase) rescue was not dependent on its TBS. This validates that the TBS requirement is due to tankyrase and further shows that C19orf43 (at least under conditions of overexpression in the absence of tankyrase) can resolve persistent telomere cohesion, independent of its TBS. Altogether, these data indicate that C19orf43 is an essential component of the persistent telomere cohesion phenotype.

### C19orf43 protects TERRA levels and suppresses R-Loops

We next asked if depleting C19orf43 would impact TERRA levels. Purified C19orf43 was shown to have exoribonuclease activity in vitro on a broad range of substrates(Xie et al., 2017). One prediction is that loss of a nuclease activity might lead to increased TERRA levels. We evaluated TERRA levels in C19orf43 KO cell lines #2 and #4 by RNA dot blot analysis. Blots were probed with ^32^P-labeled (CCCTAA)_4_ Telo C probe that hybridizes to TERRA and normalized to an 18S rRNA probe. As shown in Fig. 3A, B, we observed a significant reduction in TERRA levels. Additionally, we measured TERRA levels by Northern blot analysis of C19orf43 shRNA lines #1 and #4. Blots were probed using the Telo C probe and normalized to 18S rRNA. As shown in Fig. 3C, D, we observed a reduction in TERRA. Finally, we measured TERRA using RTqPCR in C19orf43 shRNA lines #1 and #4 cell lines using a set of subtelomere (2p, 9p, 16p, and 18p) and telomere (telo) specific primers, normalizing to the reference gene GAPDH. As shown in Fig. 3E, we observed a reduction in TERRA levels. Together these data indicate that TERRA levels are reduced in C19orf43-depleted cells.

**Figure 3.**
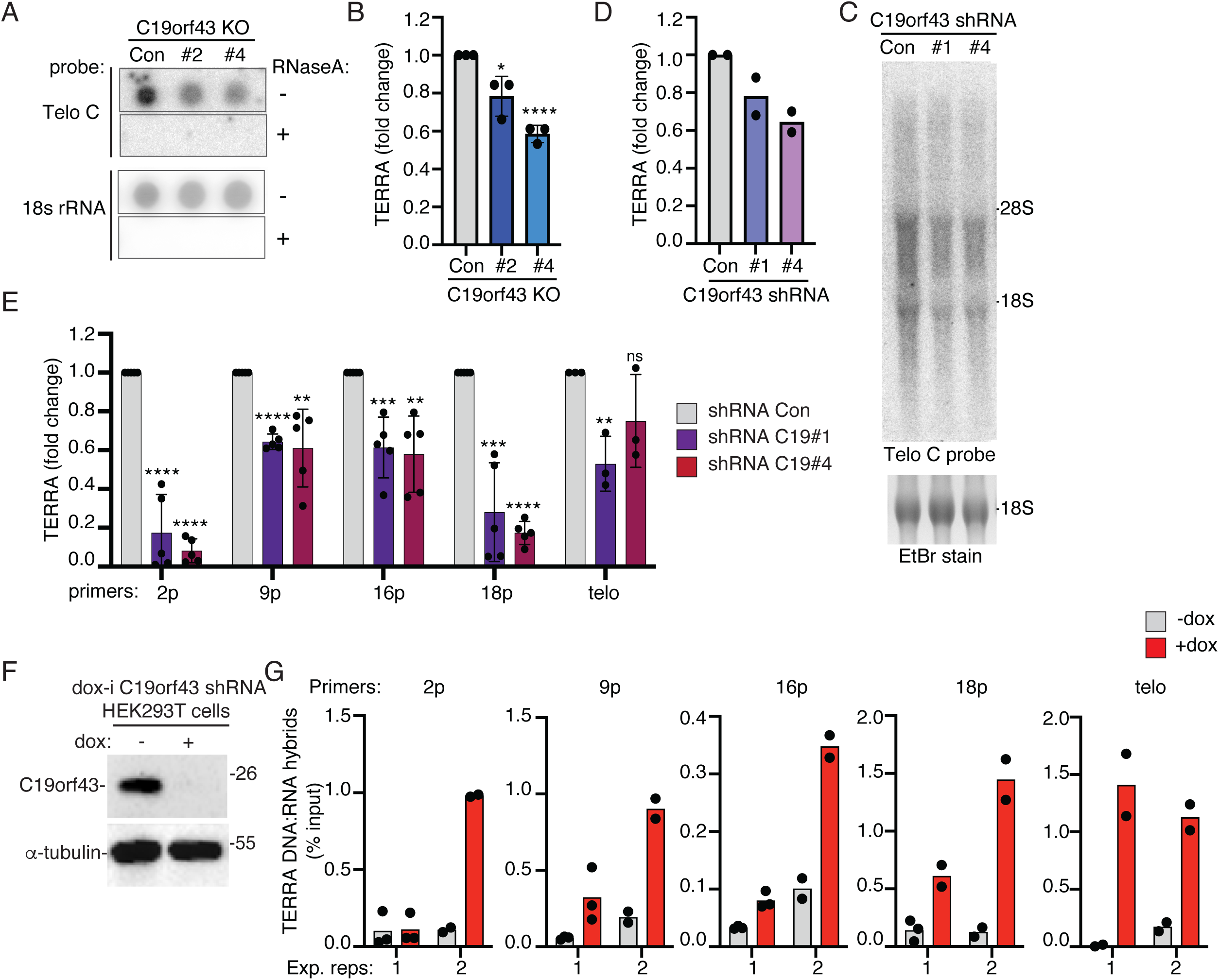
C19orf43 depletion leads to a decrease in TERRA and an increase in TERRA R-Loops. (A) RNA dot blot analysis of TERRA or 18S rRNA levels in control or C19orf43 KO#2 and #4 HEK293T cell lines. Samples were treated – or + RNaseA as a control. Blots were probed with a Telo C TERRA or 18s rRNA probe. (B) Quantification of TERRA levels normalized to 18S rRNA levels plotted as fold change over control cells. Three independent experiments ± SEM. *p :: 0.05, ****p :: 0.0001, Student’s unpaired t test. (C) Northern blot analysis of TERRA levels in control or C19orf43 shRNA #1 and #4 HEK293T cell lines. Blots were probed with a Telo C TERRA probe. (D) Quantification of TERRA levels normalized to 18S rRNA levels plotted as fold change over control cells. Two independent experiments. (E) RT-qPCR analysis of RNA samples from control or C19orf43 shRNA #1 and #4 HEK293T cell lines. TERRA levels from individual sub telomeres (2p, 9p, 16p, 18p) and all telomeres (telo probe) were measured from total RNA, normalized to GAPDH and plotted as fold change over control cells. The bars represent the average value from three to five independent experiments with two technical replicates each. ± SEM. **p :: 0.01, ***p:: 0.001, ****p :: 0.0001, Student’s unpaired t test. ns; not significant. (F) Immunoblot analysis of dox-inducible (dox-i) C19orf43 shRNA HEK293T cells following 48 hr of treatment (–) or (+) dox. (G) DRIP analysis using S9.6 antibody in extracts from -dox and +dox dox-i C19orf43 HEK293T shRNA cells. Immunoprecipitates and input samples were analyzed by qPCR with primer sets amplifying 2p, 9p, 16p, 18p subtelomeric DNA or telomeric DNA and plotted as % input. Two independent experiments (1, 2) with two to three technical replicates each.

TERRA is also found as R-loops, triple-stranded structures that comprise an RNA:DNA hybrid and a DNA single strand. We thus asked if loss of C19orf43 would impact TERRA R-loop levels. For this, we created a doxycyline-inducible lentiviral shRNA plasmid (dox-i C19orf43 shRNA) using the C19orf43 shRNA #1 sequence described above and generated a stable dox-i cell line in HEK293T cells. Cells were induced with dox for 48 hr and analyzed by immunoblot. As shown in Fig. 3F, C19orf43 levels were significantly reduced. RNA:DNA hybrid levels were assayed using DNA:RNA immunoprecipitation (DRIP) with the S9.6 antibody, which detects R-loops(Boguslawski et al., 1986). As a control, prior to immunoprecipitation, the nucleic acids were treated in situ with or without ribonuclease RNaseH1, which hydrolyses the RNA of RNA:DNA hybrids(Cerritelli and Crouch, 2009). The DNA component of the immunoprecipitated hybrid and input samples was then quantified by qPCR with primer sets amplifying 2p, 9p, 16p, 18p subtelomeric DNA or telomeric DNA and plotted as % input. As shown in Fig. 3G, we observed an increase in TERRA R-loops in the +dox (C19orf43-depleted) compared to the -dox (control) cells.

### RNaseH1 resolves persistent telomere cohesion

Our data above, indicating that depletion of C19orf43 leads to persistent telomere cohesion and to an increase in TERRA R-loops, raised the possibility that increased TERRA R-loop levels account for the persistent telomere cohesion phenotype. We thus asked if overexpression of RNaseH1 impacts persistent telomere cohesion. C19orf43 shRNA #1 cells were transfected with a vector control, RNaseH1.WT, or RNaseH1.CD (a catalytically dead mutant with a D145 mutation) (Arora et al., 2014), analyzed by immunoblot (Fig. 4A), and subjected to FISH analysis. As shown in Fig. 4B, C, we observed a robust rescue of persistent telomere cohesion with RNaseH1 WT, but not CD. We next asked if the persistent cohesion observed in TNKS DKO cells was due to R-loops. TNKS DKO cells were transfected with a vector control, RNaseH1.WT, or RNaseH1.CD, analyzed by immunoblot (Fig. 4D), and subjected to FISH analysis. As shown in Fig. 4E, F, RNaseH1 WT, but not CD rescued the persistent telomere cohesion.

**Figure 4.**
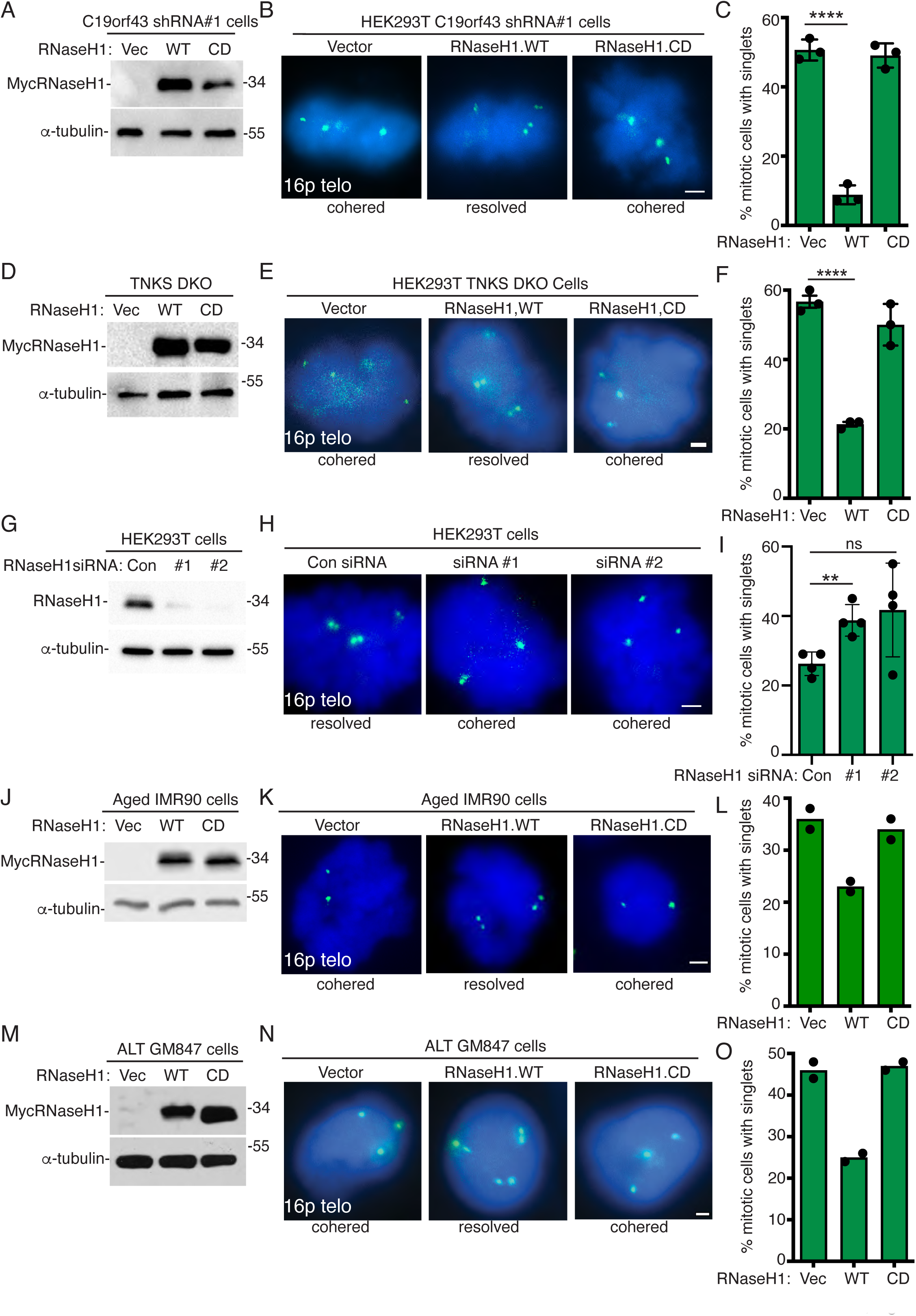
RNaseH1 overexpression resolves persistent telomere cohesion in TNKS DKO, ALT, and aged IMR90 cells. (A) Immunoblot analysis of Vector, RNaseH1.WT, or RNaseH1.CD transfected C19orf43 shRNA#1 HEK293T cell extracts. (B) FISH analysis of Vector, RNaseH1.WT, or RNaseH1.CD transfected C19orf43 shRNA#1 HEK293T mitotic cells using a 16p subtelo probe (green). (C) Quantification of the frequency of mitotic cells with cohered telomeres. Three independent experiments (n=50 cells each) ± SEM. ****p :: 0.0001, Student’s unpaired t test. (D) Immunoblot analysis of Vector, RNaseH1.WT, or RNaseH1.CD transfected HEK293T TNKS DKO cell extracts. (E) FISH analysis of Vector, RNaseH1.WT, or RNaseH1.CD transfected HEK293T TNKS DKO mitotic cells using a 16p subtelo probe (green). (F) Quantification of the frequency of mitotic cells with cohered telomeres. Three independent experiments (n=50 cells each) ± SEM. ****p :: 0.0001, Student’s unpaired t test. (G) Immunoblot analysis of Control, RNaseH1#1, or RNaseH1#2 siRNA transfected HEK293T cell extracts. (H) FISH analysis of Control, RNaseH1#1, or RNaseH1#2 siRNA transfected HEK293T mitotic cells using a 16p subtelo probe (green). (I) Quantification of the frequency of mitotic cells with cohered telomeres. Three independent experiments (n=49-87 cells each) ± SEM. **p :: 0.01, Student’s unpaired t test. ns; not significant. (J) Immunoblot analysis of Vector, RNaseH1.WT, or RNaseH1.CD transfected aged IMR90 cell extracts. (K) FISH analysis of Vector, RNaseH1.WT, or RNaseH1.CD transfected aged IMR90 mitotic cells using a 16p subtelo probe (green). (L) Quantification of the frequency of mitotic cells with cohered telomeres. Two independent experiments (n=50 cells each). (M) Immunoblot analysis of Vector, RNaseH1.WT, or RNaseH1.CD transfected ALT GM847 cell extracts. (N) FISH analysis of Vector, RNaseH1.WT, or RNaseH1.CD transfected ALT GM847 mitotic cells using a 16p subtelo probe (green). (O) Quantification of the frequency of mitotic cells with cohered telomeres. Two independent experiments (n=50 cells each). (B, E, H, K, N) DNA was stained with DAPI (blue). Scale bars represent 2 μm.

These results showed that overexpression of RNaseH1 in C19orf43-depleted or TNKS DKO cells rescued persistent telomere cohesion. To determine if RNaseH1 activity is important to resolve cohesion in wild type cells we measured the impact of RNaseH1 depletion in HEK293T cells. We introduced two distinct RNaseH1 siRNAs (#1 and #2). Immunoblot analysis showed efficient depletion of RNaseH1 (Fig. 4G) and FISH analysis showed induction of persistent telomere cohesion in the RNaseH1-depleted cells (Fig. 4H, I).

Finally, we assessed the impact of RNaseH1 in cells with naturally occurring persistent cohesion: normal aged cells and ALT cancer cells. We transfected presenescent IMR90 cells with a Vector control, RNaseH1.WT, or RNaseH1.CD, performed immunoblot analysis (Fig. 4J), and subjected cells to FISH analysis. As shown in Fig. 4K, L, persistent telomere cohesion in aged IMR90 cells was rescued by overexpression of RNaseH1.WT, but not RNaseH1.CD. Similar results were obtained with transfection into ALT GM847 cells; RNaseH1.WT (but not CD) reduced persistent telomere cohesion (Fig. 4M, N, O).

### Telomeres are cohered in mitotic cells through RNA:DNA hybrids

As described above we observed a robust rescue of persistent telomere cohesion by overexpressing RNaseH1 in cells. In these experiments RNaseH1 was transfected into C19orf43-depleted cells for 24 hrs prior to harvesting mitotic cells and subjecting them to FISH analysis. We next queried whether RNaseH1 could resolve telomere cohesion if the purified enzyme was added directly to the cohered telomeres in situ. For this approach, untransfected C19orf43-depleted mitotic cells were harvested, fixed and cytospun onto coverslips. The coverslips were then incubated with or without RNaseH1 enzyme and then analyzed by FISH (see schematized process in Fig. 5A). As shown in Fig. 5B, C, treatment with RNaseH1 in situ resolved persistent telomere cohesion.

**Figure 5.**
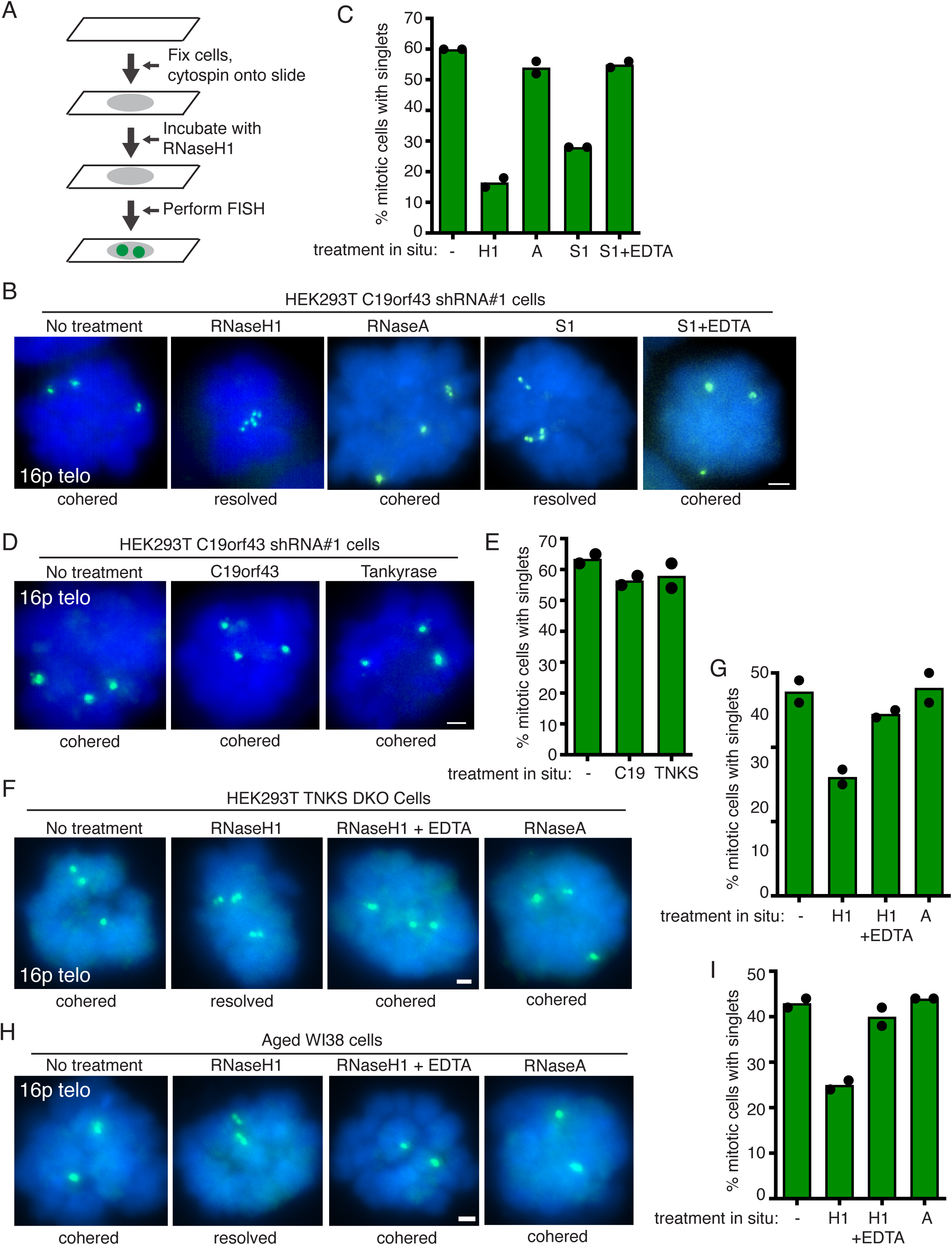
RNaseH1 resolves cohered sister telomeres in situ. (A) Schematic of experimental approach for enzyme incubations in situ prior to FISH analysis. (B) FISH analysis of C19orf43 shRNA#1 HEK293T mitotic cells following incubation in situ with no treatment, RNaseH1, RNaseA, S1, or S1 plus EDTA inhibitor, using a 16p subtelo probe (green). (C) Quantification of the frequency of mitotic cells with cohered telomeres. Two independent experiments (n=27-50 cells each). (D) FISH analysis of C19orf43 shRNA#1 HEK293T mitotic cells following incubation in situ with no treatment, C19orf43, or tankyrase, using a 16p subtelo probe (green). (E) Quantification of the frequency of mitotic cells with cohered telomeres. Two independent experiments (n=40-50 cells each). (F) FISH analysis of TNKS DKO HEK293T mitotic cells following incubation in situ with no treatment, RNaseH1, RNaseH1 plus EDTA inhibitor, or RNaseA using a 16p subtelo probe (green). (G) Quantification of the frequency of mitotic cells with cohered telomeres. Two independent experiments (n=50 cells each). (H) FISH analysis of aged WI38 mitotic cells following incubation in situ with no treatment, RNaseH1, RNaseH1 plus EDTA inhibitor, or RNaseA using a 16p subtelo probe (green). (I) Quantification of the frequency of mitotic cells with cohered telomeres. Two independent experiments (n=50 cells each). (B, D, F, H,) DNA was stained with DAPI (blue). Scale bars represent 2 μm.

Our in situ analysis indicated that the cohered telomeres (singlets) were held together by RNA:DNA hybrids. We sought to further query the nature of the cohered telomeres by incubating the coverslips with other enzymes in situ. We tested RNaseA, an endoribonuclease that specifically degrades single-stranded RNA and not the RNA:DNA hybrid(Laspata et al., 2023; Zhang et al., 2015). As shown in Fig. 5B, C, incubation with RNaseA in situ did not resolve cohered telomeres. We next tested S1, an endonuclease that cleaves single stranded DNA. As shown in Fig. 5B, C, sister telomere cohesion was resolved by S1. Resolution was blocked by inclusion of the chelator EDTA, which inhibits S1. Together, the in situ analysis indicates that persistent telomere cohesion is mediated by RNA:DNA hybrids and single-stranded DNA, but not single-stranded RNA.

We additionally incubated fixed mitotic cells with purified recombinant C19orf43 protein or recombinant tankyrase in situ. However, as shown in Fig. 5D, E unlike RNaseH1, neither C19orf43 nor tankyrase resolved persistent telomere cohesion in situ. Thus, while C19orf43 and tankyrase (like RNaseH1) promote resolution of telomere cohesion when expressed inside cells, (unlike RNaseH1) they cannot promote resolution of the cohered sisters when incubated with fixed mitotic cells in situ.

We next asked if the cohered telomeres in TNKS DKO mitotic cells could be resolved by RNaseH1 in situ. TNKS DKO mitotic cells were harvested, fixed, cytospun onto coverslips, incubated with or without RNaseH1 enzyme, and then analyzed by FISH. As shown in Fig. 5F, G, treatment with RNaseH1 in situ resolved persistent telomere cohesion. Resolution was blocked by inclusion of the chelator EDTA, which inhibits RNaseH1. Incubation with RNaseA in situ did not resolve cohered telomeres (Fig. 5F, G). Similar results were obtained when the analysis was performed on aged WI38 cells. Treatment with RNaseH1, but not RNaseA, of aged WI38 mitotic cells in situ led to resolution of persistent telomere cohesion (Fig. H, I). Together our data indicate that persistent telomere cohesion in C19orf43-depleted, TNKS DKO, and aged WI38 cells is mediated by RNA:DNA hybrids.

### C19orf43 depletion extends replicative lifespan of aged IMR90 cells

Persistent telomere cohesion occurs naturally in normal human cells as they approach senescence. We showed previously that this has a protective role; forced resolution of cohesion in aged cells by overexpression of TRF1 led to DNA damage and a senescent-like growth arrest(Azarm et al., 2020). We wondered if inducing persistent cohesion in young cells (at an early PD) might confer benefits as the cells aged. For this, we generated a stable dox-i C19orf43 shRNA IMR90 cell line at an early PD (PD25). Cells were induced with dox for 48 hr and analyzed by immunoblot. As shown in Fig. 6A, C19orf43 levels were significantly reduced in the +dox cells. FISH analysis showed induction of persistent telomere cohesion in the +dox (C19orf43-depleted) cells (Fig. 6B, C). The cells were then passaged in culture –dox or +dox for 120 days. Initially (for the first 40 days) the cells grew at the same rate with or without dox (Fig. 6D). However, after 40 days (PD37) the growth rate of the +dox (C19orf43-depleted) cells increased compared to the -dox (control) cells. The -dox (control) cells growth rate began to slow and ultimately ceased at 65 days (PD48). The +dox (C19orf43-depleted) cells continued to divide for an additional 55 days (PD62) (Fig. 6D). Thus, depletion of C19orf43 decidedly extended the replicative lifespan of the aged cells.

**Figure 6.**
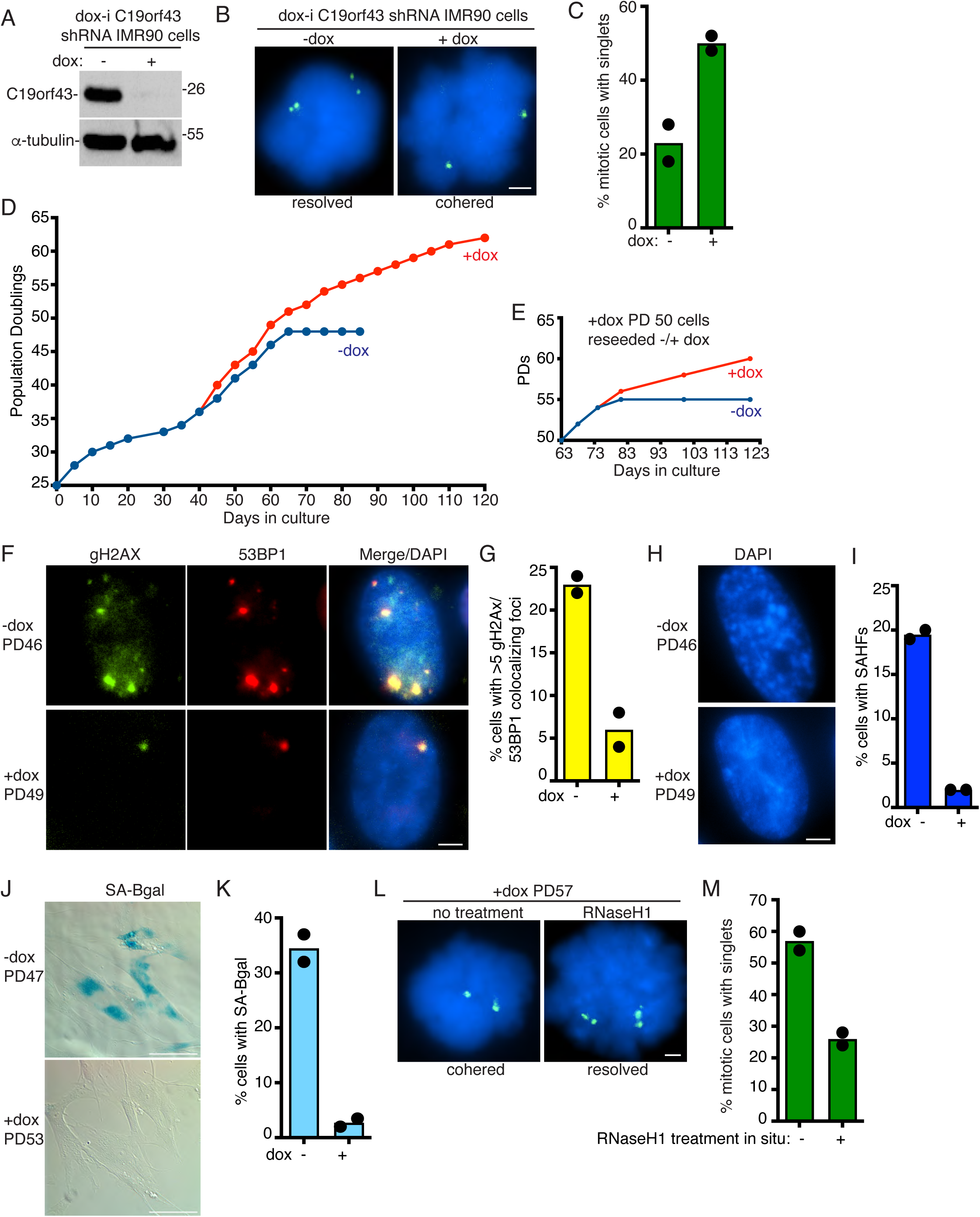
C19orf43 depletion delays replicative senescence. (A) Immunoblot analysis of dox-i C19orf43 shRNA IMR90 cells (-dox PD43) following 48 hrs (-) or (+) dox. (B) FISH analysis of mitotic dox-i C19orf43 shRNA IMR90 cells (-dox PD39) following 48 hrs (-) or (+) dox using a 16p subtelo probe (green). (C) Quantification of the frequency of mitotic cells with cohered telomeres. Two independent experiments (n=50 cells each). (D) Growth curve analysis of dox-i C19orf43 shRNA IMR90 (PD25) cells generated by lentiviral infection and passaged (-) or (+) dox for 120 days. (E) Growth curve analysis of dox-i C19orf43 shRNA IMR90 (+dox PD50) cells reseeded (–) or (+) dox and passaged for an additional 60 days. (F) Immunofluorescence analysis of dox-i C19orf43 shRNA IMR90 -dox PD46 or +dox PD49 cells stained with γH2AX (green) and 53BP1 (red) antibodies. (G) Quantification of the frequency of cells displaying >5 γH2AX/53BP1 colocalizing foci from -dox PD43/46 or +dox PD45/49 cells. Two independent experiments (n=106-124 cells each). (H) Detection of senescence-associated heterochromatin foci (SAHF) in DAPI-stained dox-i C19orf43 shRNA IMR90 -dox PD46 or +dox PD49 cells. (I) Quantification of SAHF-positive DAPI-stained dox-i C19orf43 shRNA IMR90 -dox PD43/46 or +dox PD45/49 cells. Two independent experiments (n=182-215 cells each). (J) SA-β-gal analysis of dox-i C19orf43 shRNA IMR90 -dox PD47 or +dox PD53 cells. Scale bars represent 100 μm. (K) Quantification of SA-β-gal positive cells. Two independent experiments (n=172-207 cells each). (L) FISH analysis using a 16p subtelo probe (green) of mitotic dox-i C19orf43 shRNA IMR90 +dox (PD57) cells following no treatment or RNaseH1 treatment in situ. (M) Quantification of the frequency of mitotic cells with cohered telomeres. Two independent experiments (n=50 cells each). (B, F, H, L) DNA was stained with DAPI (blue). Scale bars represent 2 μm.

To determine if the growth advantage conferred by C19orf43 depletion was reversible, PD50 +dox (C19orf43-depleted) cells were reseeded with or without dox (Fig. 6E). Initially (for the first 10 days) the cells grew at the same rate with or without dox. However, after 10 days the growth rate of the +dox (C19orf43-depleted) cells increased compared to that of the -dox (control) cells. The -dox (control) cells began to slow down and ultimately ceased growth at 81 days (PD55) while the +dox (C19orf43-depleted) cells continued to divide (Fig. 6E).

Aged human cells accumulate DNA damage that signals senescence(d’Adda di Fagagna et al., 2003). We thus asked if the DNA damage signal was attenuated in the +dox (C19orf43-depleted) versus the -dox (control) cells by measuring the frequency of DNA damage foci at late PDs (∼day 60). As shown in Fig. 6F, G, -dox (control) (PD46) cells displayed a high level of DNA damage foci that was dramatically reduced in the +dox (C19orf43-depleted) (PD49) cells. To determine if senescence was delayed in the +dox (C19orf43-depleted) versus the -dox (control) cells we measured senescence associated heterochromatin foci (SAHF)(Narita et al., 2003) in late PD cells and observed a dramatic reduction in SAHFs in +dox (C19orf43-depleted) versus -dox (control) cells (Fig. 6H, I,). Finally, measurement of the senescence associated marker β-galactosidase (SA-β-gal)(Dimri et al., 1995) revealed a dramatic reduction in SA-β-gal-positive cells in +dox (C19orf43-depleted) (PD53) versus -dox (control) (PD47) cells, indicating delayed senescence (Fig. 6J, K).

To determine the status and nature of the cohesion in cells with extended lifespan, we performed FISH analysis on the +dox (C19orf43-depleted) cells at PD57 (day 90). Prior to FISH, we incubated the cells in situ without or with RNaseH1. As shown in Fig. 6L, M, the +dox (C19orf43-depleted) cells (without treatment) exhibited a high level of persistent telomere cohesion. Incubation of the coverslips with RNaseH1 in situ led to resolution of cohesion. Together these data indicate that aged C19orf43-depleted cells exhibit cohered telomeres in mitosis that are held together by RNA:DNA hybrids and are protected from DNA damage and senescence.

## Discussion

Telomere shortening robs normal human cells of their potential for immortality and thus serves as a roadblock to tumorigenesis. Chromosome replication in the absence of telomerase erodes the reservoirs of TTAGGG repeats, as well as the proteins that bind them. Ultimately, this loss culminates in cessation of growth. This does not occur suddenly, but rather through the gradual process known as replicative senescence. We describe a mechanism inherent to the loss of telomere repeats (and associated proteins) that shepherds chromosome ends through this process. Integrity of the genome relies upon mechanisms that prevent accumulation of potentially toxic structures such as R-loops. Shortened telomeres lose their ability to prevent R-loop buildup. Surprisingly, these otherwise toxic structures afford protection to critically short telomeres, promoting a gradual replicative senescence.

Persistent telomere cohesion in mitosis is a state that occurs naturally in cells that lack telomerase: normal aged cells and ALT cancer cells. This state can also be created artificially in telomerase positive cells by depletion of tankyrase, mutation of the tankyrase binding site in TRF1, and now in this report, by depletion of C19orf43 or mutation of its tankyrase binding site. We suggest a model that when telomeres are sufficiently long, C19orf43 is recruited and binds TERRA to limit R-loop formation. Upon telomere shortening, the TRF1/tankyrase/C19orf43 axis is diminished leading to R-loop accumulation and persistent telomere cohesion. Unexpectedly, the increase in TERRA R-loops was accompanied by a decrease in TERRA levels. This was surprising because in other examples increased R-loops was accompanied by increased TERRA levels(Arora et al., 2014; Graf et al., 2017; Sagie et al., 2017). There are likely multiple ways to regulate R-loops at telomeres. In the case of C19orf43 depletion, the increased R-loops could block TERRA transcription, leading to a reduction in TERRA levels. Interestingly, a recent study used a proximity labeling-based approach to identify the RNase H proximal proteome and found that C19orf43 (among a number of other proteins) was significantly enriched(Yan et al., 2022). The study suggested that there were multiple classes of R-loop regulators that use distinct mechanisms. C19orf43 could be part of an R-loop repressing cellular machinery that through its association with TRF1/tankyrase limits R-loops at long, but not critically short, telomeres.

What actually holds persistently cohered sister telomeres together? We showed that overexpression of RHaseH1 in living cells resolves persistent telomere cohesion, indicating a role for RNA:DNA hybrids. Conversely and in support, depletion of RNaseH1 led to increased persistent telomere cohesion. Remarkably, the action of RNaseH1 could be recapitulated in situ; incubation of fixed cohered telomeres with purified RNaseH1 in situ resolved telomere cohesion. Thus, RNA:DNA hybrids hold sisters together. R-loops are comprised of RNA:DNA hybrids and a displaced single strand. The displaced DNA single strand could invade the sister telomere and connect sisters. In support of this, treatment of cohered telomeres in situ with the S1 endonuclease, which cleaves single stranded DNA, promoted resolution of cohesion. We suggest that persistent cohesion reflects TERRA R-loop mediated associations between sister telomeres. These associations could protect critically short telomeres from being viewed as DNA breaks in G2/M. The question remains how these associations would ultimately get resolved in mitotic cells. We showed previously by live cell imaging that persistent telomere cohesion (in tankyrase depleted or aged IMR90 cells) led to anaphase delay(Kim and Smith, 2014). Cells struggled briefly to segregate their chromosomes, but ultimately resolved them and proceeded through the cell cycle without damaged telomeres. The R-loop mediated interactions described here could (with the aid of spindle forces in live cells) enable resolution in anaphase without significant damage.

When young IMR90 cells were depleted of C19orf43 (+dox) and passaged in culture they grew for 55 days (14 PDs) more than their uninduced -dox counterpart. When +dox (C19orf43-depleted) cells at PD50 were reseeded +/– dox, the -dox (control) cells rapidly ceased growth. Thus, the persistent cohesion (R-loop connection) affords only a transient protection. Ultimately, even the +dox (C19orf43-depleted) cells succumb to senescence. We showed previously that at the senescence point (the final PD) sister telomere cohesion is lost, likely due to the inability of short telomeres to establish cohesion(Azarm et al., 2020). Thus, persistent telomere cohesion offers a transient protective state, but cannot ultimately override telomere loss. Nonetheless, delaying or prolonging senescence could impact organismal fitness. While replicative senescence is an essential barrier to tumorigenesis, accumulation of senescent cells can impact aging and age-related diseases including cancer(Campisi, 2013; Rossiello et al., 2022). Thus, the ability to modulate/lengthen the senescence window could offer therapeutic opportunities.

## Materials and Methods

### Plasmids

The shRNA plasmids against C19orf43 were generated by cloning hairpins targeting the following sequences into the pLKO.1 puro vector: C19orf43#1 against the 3’ UTR 5’- GCCTCGTGAGACTTCATAGAA-3’ and C19orf43#4 against the coding sequence 5’- AGACGGAGGATGAGGTATTAA-3’. For dox-i C19orf43 the same hairpin as for #1, the 3’ UTR 5’- GCCTCGTGAGACTTCATAGAA-3’, was cloned into pLKO-Tet-On (Addgene #21915).

RNaseH1 plasmids comprised of C-terminally myc-tagged human full length RNaseH1, wild type (WT) or catalytically dead (CD) (D145A), cloned into the retroviral vector pLHCX (Clontech) were kindly provided by Claus Azzalin(Arora et al., 2014).

The C19orf43.WT plasmid is comprised of an N-terminal 3x Flag tag followed by human full length C19orf43 cloned into the p3XFlag-CMV-10 vector (Sigma). C19orf43.R7G was obtained by mutating the arginine at position 7 to glycine and C19orf43.R106G was obtained by mutating the arginine at position 106 to glycine. Mutagenesis was performed using Q5 Site-Directed Mutagenesis Kit (NEB) according to the manufacturer’s instructions. For expression in E. coli, C19orf43 was cloned into the pET-22b vector with a C-terminal His tag and expressed without the pelB leader sequence.

### Cell lines

WI38 (ATCC) and IMR90 (ATCC) fibroblast cell lines were supplemented with 20% FBS and grown in standard conditions. GM847 (ATCC) ALT cells, HTC75 (van Steensel and de Lange, 1997), HEK293T (ATCC) and TNKS1/2 DKO(Bhardwaj et al., 2017) cell lines were supplemented with 10% FBS and grown in standard conditions.

### Generation of C19orf43 tankyrase binding site (TBS) mutant and knockout (KO) cell lines using CRISPR/Cas9

C19orf43 TBS mutant and KO cell lines were generated using RNA-guided CRISPR associated nuclease Cas9. A 20 bp target sequence directed against the first exon of the human C19orf43 gene (C19 guide DNA 5’–GACGGGCGGAGCCTCAGGGC–3′) was inserted into the guide sequence insertion site using Bbs1 site of the CRISPR plasmid pX330 comprised of Cas9 and a chimeric guide RNA(Shalem et al., 2014). To generate C19orf43 TBM clones, a single stranded DNA homology template was designed to change Gly12 to Ala (C19orf43.G12A) and also to introduce an XhoI restriction site to screen the potential clones. To generate the C19orf43 KO clones, a single stranded DNA homology template was designed to change Arg6 to a stop codon (TAA) and also to introduce an XhoI restriction site to screen the potential clones.

HEK293T cells were transfected with the pX330 plasmid and homology template ssDNA as described previously(Ran et al., 2013). Following transfection, cells were re-plated for single cell cloning, propagated and screened by a PCR strategy designed to screen for gain of an XhoI site in the target site, using the forward primer Fwd 5’ – CTTCCCGGCATGCATTGTTC -3′ or Fwd1 5′- AACCCGCGAGACGGGGGCT-3′ and the reverse primer Rev 5′-CACGGCTCCTTACGAAGCTA-3′.

Three independent homozygous C19orf.G12A clones (#3, #4, and #6) were isolated and confirmed by DNA sequencing of the PCR products. Two independent homozygous KO clones (#2 and #4) were isolated and confirmed by DNA sequencing of the PCR products.

### siRNA and plasmid transfection

For plasmids, cells were transfected with Lipofectamine 3000 (Invitrogen) according to the manufacturer’s protocol for 20 hr.

For siRNAs, cells were transfected with Lipofectamine RNAiMAX (Invitrogen) according to the manufacturer’s protocol for 48-72 hr. The final concentration of siRNA was 20 nM. The following target sequences were used for the RNaseH1 siRNAs: RNaseH1#1 (5’- TCCTTTAAATGTAGGCATTAGACTT-3’) described previously as RNaseH1c(Arora et al., 2014), and RNaseH1#2 (5’-GGGAAAGAGGTGATCAACA-3’) described previously as RNAseH1-2 siRNA(Yadav et al., 2022) (Dharmacon). The Control is the GFP duplex I (Dharmacon).

### Lentiviral Infection

shRNA cell lines were generated by introducing lentiviruses expressing GFP (Control), C19orf43#1, C19orf43#4 or dox-i C19orf43 into HEK293T or IMR90 cells. For lentivirus generation, 293FT cells (Invitrogen) were transfected using Lipofectamine 3000 (Invitrogen) with 1 μg each lentiviral vector and pCMVΔR8.9 packaging plasmid, and 100 ng pMD.G envelope plasmid. Forty-eight hours after transfection, supernatants were collected, filtered with a 0.45- μm filter (Millipore), supplemented with 8 μg/ml polybrene (Sigma-Aldrich), and used to infect target cells. Following 48–72 h infection, cells were sub-cultured 1:2 into medium containing 2 μg/ml puromycin.

### Preparation of cell extracts

Cells were resuspended in four volumes of TNE buffer [10 mM Tris (pH 7.8), 1% Nonidet P-40, 0.15 M NaCl, 1 mM EDTA, and 2.5% protease inhibitor cocktail (PIC) (Sigma)] and incubated for 1 hr on ice. Suspensions were pelleted at 10,000 x g for 10 min at 4C. Equal amounts of supernatant proteins (determined by Bio-Rad protein assay) were fractionated by SDS-PAGE and analyzed by immunoblotting.

### Immunoblot analysis

Immunoblots were incubated separately with the following primary antibodies: mouse anti-Myc 05-724 (0.1 µg/ml; Millipore), mouse anti-Flag F3165 (3.8 µg/ml, Sigma), rabbit anti-C19orf43 PA5-63805 (0.4 µg/ml, Invitrogen), rabbit anti-TNKS1 762 (1 µg/ml) (Scherthan et al., 2000), rabbit anti-RNaseH1 15606-AP (0.5 µg/ml, Proteintech), or mouse anti-α-tubulin ascites (1:10000-1:20000, Sigma), followed by horseradish peroxidase-conjugated donkey anti-rabbit or anti-mouse IgG (1:3000, Amersham). Bound antibody was detected with Super Signal West Pico (Thermo Scientific).

### Immunoprecipitation

Cells were lysed as above and supernatants precleared with Protein G-Sepharose rotating at 4 °C for 30 min. Nonspecific protein aggregates were removed by centrifugation and the supernatant was used for immunoprecipitation analysis or fractionated directly on SDS-PAGE (indicated as input, ∼5% of the amount used in the immunoprecipitation). For immunoprecipitation of Flag epitope-tagged proteins, supernatants were incubated with 20 μl of Flag M2 agarose (Sigma, A2220) for 3 h. For TNKS immunoprecipitations supernatants were incubated for 3 h with 1 μg rabbit anti-tankyrase 465(Smith et al., 1998) or IgG, followed by Protein G Sepharose for 1 h. For all immunoprecipitations, beads were washed three times with 1ml of TNE buffer, fractionated by SDS-PAGE, and processed for immunoblotting as described above.

### Chromosome-specific FISH

Cells were fixed and processed as described previously (Dynek and Smith, 2004). Briefly, cells were isolated mechanically by mitotic shake-off, fixed twice in methanol:acetic acid (3:1) for 15 min, cytospun (Shandon Cytospin) at 2,000 rpm for 2 min onto slides, rehydrated in 2X SSC at 37°C for 2 min, and dehydrated in an ethanol series of 70%, 85%, and 100% for 2 min each. Cells were denatured at 75°C for 2 min and hybridized overnight at 37°C with FITC-conjugated 16ptelo subtelomere probe or FITC-conjugated subtelomere and TRITC-conjugated arm 13q14.3 deletion probe (13qtelo/13qarm) from Cytocell. Cells were washed in 0.4X SSC at 72°C for 2 min, and in 2X SSC with 0.05% Tween 20 at RT for 30 seconds. DNA was stained with 0.2 µg/ml DAPI. For FISH analysis of WI38 and IMR90 fibroblasts, cells were treated with 50 ng/ml nocodazole (Sigma) for 16 hr prior to shake-off. Mitotic cells were scored as having telomeres cohered (singlets) if 50% or more of their loci appeared as singlets, i.e., one out of two or two out of three.

For in situ treatment with purified enzymes, following cytospin samples were rehydrated in 50 µl and subjected to the following treatments. Coverslips were incubated with: 5 units RNaseH1 (NEB) in RNaseH1 Buffer (50 mM Tris-HCl pH 8.3, 75 mM KCl, 3 mM MgCl_2_,10 mM DTT) and incubated for 30 to 60 min without or with 6 mM EDTA at 37°C; RNaseA (0.2mg/ml final concentration) (Invitrogen) in H_2_0 for 30 to 60 min at 37°C; 10 units S1 Nuclease (Invitrogen) in S1 Buffer (30 mM sodium acetate pH 4.6, 50 mM NaCl, 1 mM zinc acetate, and 5% glycerol) without or with 30 mM EDTA (pretreated for 10 min at 70C) for 30 min at 37°C; 6 µg C19orf43 (purified from E. coli BL21 through Qiagen Ni-TA agarose using standard procedures) in RNaseH1 buffer for 30 min at RT; or with 2 µg tankyrase (purified from baculovirus as described previously) (Smith et al., 1998) with 100 mM NAD^+^ in 50 mM Tris pH 8.0, 4mM MgCl2, 0.2 mM DTT for 30 min at RT. Following treatment cells were washed 2X with 2X SSC and processed as described above beginning with the 2X SSC treatment at 37°C for 2 min.

### Indirect Immunofluorescence

Cells were fixed in 2% paraformaldehyde in PBS for 10 min at RT, permeabilized in 0.5% NP-40/PBS for 10 min at RT, blocked in 1% BSA/PBS, and incubated with mouse anti-γH2AX #05 636 (20 µg/ml; Millipore) and rabbit anti-53BP1 NB 100-304 (1 µg/ml; Novus Biologicals). Primary antibodies were incubated at RT for 2 hr, followed by detection with FITC-conjugated or TRITC-conjugated donkey anti-rabbit or anti-mouse antibodies (1:100; Jackson Laboratories). DNA was stained with 0.2 µg/ml DAPI.

### Detection of Senescence-associated heterochromatin foci (SAHF)

SAHF was analyzed on coverslips processed for γH2AX and 53BP1 immunofluorescence described above. A cell was scored as SAHF-positive if its DAPI counterstain had a characteristic punctate pattern (Narita et al., 2003).

### Senescence associated β-galactosidase assay

For the SA-β-galactosidase assay (Dimri et al., 1995), cells were fixed in 2% formaldehyde and 0.2% glutaraldehyde in PBS for 5 min, washed three times in PBS, and stained for 5 hr at 37°C in staining solution (1 mg/mL X-gal, 150 mmol/L NaCl, 2 mmol/L MgCl2, 5 mmol/L K3Fe[CN]6, 5 mmol/L K4Fe[CN]6, and 40 mmol/L NaPi, pH 6.0).

### RNA isolation and Dot Blot

RNA was isolated from cells using RNeasy Mini Kit (Qiagen). Two on-column (Qiagen) and one in-solution (NEB) DNase digestions were performed. Purified RNA was digested with RNase (DNase-free) (Invitrogen), as a control. All samples were denatured at 65°C for 5 min, cooled on ice 5 min, and blotted (10 μg per sample) onto a Hybond-XL membrane (Amersham) using a dot-blot apparatus (Bio-Rad). RNA was UV-crosslinked to the membrane. The membrane was then blocked in Church buffer (0.5 M NaHPO4, 1 mM EDTA, pH 8.0, 1% (w/v) BSA, 7% SDS) for at least 2 hr at 55°C, and then hybridized to a ^32^P end-labeled (CCCTAA)_4_ oligonucleotide Telo C probe in Church Buffer o/n 55°C. The membrane was washed 3-4 times, 15 minutes each with 4X SSC at RT and exposed to a phosphorimager screen. Radioactive signal was detected with a Typhoon Biomolecular Imager (GE). After signal detection, the membrane was stripped by incubation with boiling 0.1 % SDS, 2mM EDTA three times. The membrane was then blocked in Church buffer for 2 hr at 55°C, and then hybridized to a ^32^P end-labeled (CCATCCAATCGGTAGTAGCG) 18s rRNA probe at 55°C overnight and processed as describe for the Telo C probe.

### Northern Blot analysis

RNA was isolated as described above. 15 μg total RNA per sample was subjected to electrophoresis on a 1.3% agarose formaldehyde gel. The gel was stained with ethidium bromide, the RNA visualized by UV, and (capillary) transferred to Hybond-XL membrane (Amersham). The RNA was UV-crosslinked and processes as described above for the dot blot with Telo C probe.

### RTqPCR

TERRA RT-qPCR was performed as previously described with some modifications(Feretzaki and Lingner, 2017). Cells were treated with 25ng/ml nocodazole for 16 hr prior to harvest. RNA was isolated from cells using RNeasy Mini Kit (Qiagen). Two on-column (Qiagen) and one in-solution DNase (NEB) digestions were performed. Five micrograms of RNA was reverse-transcribed using 200 U SuperScript III Reverse transcriptase (Thermo Fisher Scientific), and GAPDH (5’-GCCCAATACGACCAAATCC-3’) and TERRA (5’- CCCTAACCCTAACCCTAACCCTAACCCTAA-3’) reverse primers in a 20 µl reaction. Reverse transcription was performed at 55°C for 1 h, followed by heat inactivation at 70°C for 15 min. Sample was diluted to 80 µl with H20. For each qPCR, 3 µl was mixed with Power SYBR Green PCR Master mix (Applied Biosystems) and 0.5 μM forward and reverse qPCR primers. qPCR consisted of 10 min at 95°C followed by 40 cycles at 95°C for 15 sec and 60°C for 1 min in an Applied Biosystems StepOne Real-Time System. The following primers were use: subtelomeric primers(Feretzaki et al., 2019): 2p (forward primer: 5’- GTAAAGGCGAAGCAGCATTCTCC-3’, reverse primer: 5’- TAAGCCGAAGCCTAACTCGTGTC -3’); 9p (forward primer: 5’- GAGATTCTCCCAAGGCAAG -3’, reverse primer: 5’- ACATGAGGAATGTGGGTGTTAT -3’); 16p (forward primer: 5’- TGCAACCGGGAAAGATTTTATT-3’, reverse primer: 5’- GCCTGGCTTTGGGACAACT -3’); 18p (forward primer: 5’- TACCTCGCTTTGGGACAAC -3’, reverse primer: 5’- CCTAACCCTCACCCTTCTAAC-3’); and telomeric primers(Cawthon, 2002) (forward primer: 5’- GGTTTTTGAGGGTGAGGGTGAGGGTGAGGGTGAGGGT-3’, reverse primer: 5’- TCCCGACTATCCCTATCCCTATCCCTATCCCTATCCCTA -3’).

### DRIP

DNA–RNA immunoprecipitation performed as previously described with some modifications(Feretzaki et al., 2020; Hatchi et al., 2015). Cells were treated with 25ng/ml nocodazole for 16 hr prior to harvest. Cells were harvested, counted, and washed with PBS. Up to ten million cells were dissolved in 175 μl of ice-cold RLN buffer (50 mM Tris-HCl pH 8.0, 140 mM NaCl, 1.5 mM MgCl2, 0.5% NP-40, 1 mM dithiothreitol DTT, and 100 U ml SuperaseIn (Ambion), incubated on ice for 5 min, and pelleted 300xg for 5 min 4°C. The nuclear pellet was warmed to RT and lysed in 500 μl RA1 buffer (Machery Nagel) containing 5 μl of β- mercaptoethanol, and homogenized by passing through a 20 G x 1-1/2 inch hypodermic needle (BD). The nucleic-acid-containing extracts (500 μl) were mixed with 250 μl H2O and 750 μl phenol:chloroform:isoamylalcohol (25:24:1) in a Phase Lock Gel heavy (5PRIME) 2 ml tube and pelleted (13,000xg, 5 min, RT). The upper aqueous phase was mixed with 1500 μl of ice-cold isopropanol and 30 ul 5M NaCl (to 50 mM), then incubated on ice for 30 min; precipitated nucleic acids pelleted at 10,000xg for 30 min at 4C. The pellets were washed twice with 70% ice-cold ethanol, air-dried, dissolved in 140 μl of H2O, and sonicated with Covaris R230 Focused-Ultrasonicator with R230_500750 PSU Rack 96 microTube Plate +0.5 offset (25% duty factor, 450 peak incident power, 600 cycles per burst, 70 seconds, 130 μl) to obtain fragments of 350 bp. Nucleic acid concentration was determined by nanodrop. Five μg of sonicated nucleic acids in 40 ul of 1X RNaseH1 buffer (20 mM HEPES-KOH pH 7.5, 50 mM NaCl, 10 mM MgCl2, 1 mM DTT) were treated with 3 ul (15 units) RNaseH1 (NEB) or H20 as a control and incubated at 37°C for 2 hr with slight agitation (300 rpm). Samples were diluted to 1400 ul with IP buffer (16.6 mM Tris pH 8.0, 166mM NaCl, 1.2 mM EDTA pH 8.0, 1.1% Triton X-100, 0.01% SDS) and pre-cleared with 40 μl of sepharose protein G beads (50% slurry) for 1 h, on a rotating wheel, at 4°C. 650 μl was used for each IP and one per cent (6.5 μl) of the nucleic acids were kept as input. 650 ul of sample (roughly 2.5 μg of nucleic acids) was incubated with 1 μg of S9.6 antibody (Kerafest) or mouse IgG and 30 μl of sepharose protein G beads on a rotating wheel at 4°C overnight. The next day the samples were washed for 5 min on a rotating wheel at 4°C with 1 ml each of the following buffers: A (20 mM Tris pH 8.0, 165 mM NaCl, 2 mM EDTA pH 8.0, 1% Triton X-100, 0.1% SDS), B (50 mM 20 mM Tris pH 8.0, 500 mM NaCl, 2 mM EDTA pH 8.0, 1% Triton-X100, 0.1% SDS), C (10 mM Tris-HCl pH 8.0, 1 mM EDTA pH 8.0, 250 mM LiCl, 1% NP-40, 1% Na-deoxycholate), and D (10 mM Tris-HCl pH 8.0, 1 mM EDTA pH 8.0). Beads were incubated with 50 μl IP elution buffer (100 mM NaHCO3, 1 % SDS), twice 60 min each at 65°C shaking (1050 rpm). Inputs were adjusted to 100 μl with IP elution buffer. The 100 ul samples were digested with RNaseA DNase-free (Thermofisher) 10 μg each at 37°C for 1 hr and then with 4 μg Proteinase K each at 45°C for 1 hr, and purified using GFX PCR DNA purification kit (Cytiva). DNA was eluted twice with 30 μl H20 and 1 μl used for qPCR as described above

### Image acquisition

FISH images were acquired with a Zeiss Axioplan 2 microscope with a Plan Apochrome 63X NA 1.4 oil immersion lens and a digital camera (C4742-95; Hamamatsu Photonics) using Openlab software (Perkin Elmer) or a Nikon Eclipse Ti microscope with a Nikon Plan APC TIRF 60X 1.45 oil ANDOR Neo/Zyla camera using NIS-ElementsAR4.20.02 64-bit software. For chromosome specific FISH, if foci fell in more than one optical plane of focus, multiple planes were merged using Openlab software for the Zeiss images or the Fiji application for the Nikon images. Immunofluorescence images were acquired using a Nikon Eclipse Ti microscope. SA-β-gal stained images were imaged with simple brightfield at 20X magnification using a Zeiss AxioObserver.Z1 microscope and a Axiocam 503 camera.

### Statistical analysis

Statistical analysis was performed using Prism 9 software. Data are shown as mean ± SEM. Student unpaired *t* test was applied. P < 0.05 values were considered significant: *, P :: 0.05; **, P :: 0.01; ***, P :: 0.001; ****, P:: 0.0001; ns, not significant.

## Acknowledgements

We thank Jerome Perrard for helpful discussions and critical reading of the manuscript. Research reported in this publication was supported by the National Institutes of Health under award number R01GM141292.

